# Developmental profile of psychiatric risk associated with voltage-gated cation channel activity

**DOI:** 10.1101/2020.10.19.345801

**Authors:** Nicholas E Clifton, Leonardo Collado-Torres, Emily E Burke, Antonio F Pardiñas, Janet C Harwood, Arianna Di Florio, James TR Walters, Michael J Owen, Michael C O’Donovan, Daniel R Weinberger, Peter A Holmans, Andrew E Jaffe, Jeremy Hall

## Abstract

**Background:** Recent breakthroughs in psychiatric genetics have implicated biological pathways onto which genetic risk for psychiatric disorders converges. However, these studies do not reveal the developmental time point(s) at which these pathways are relevant.

**Methods:** We aimed to determine the relationship between psychiatric risk and developmental gene expression relating to discrete biological pathways. We used post-mortem RNA sequencing data (BrainSeq and BrainSpan) from brain tissue at multiple pre- and post-natal timepoints and summary statistics from recent genome-wide association studies of schizophrenia, bipolar disorder and major depressive disorder. We prioritised gene sets for overall enrichment of association with each disorder, and then tested the relationship between the association of each of their constituent genes with their relative expression at each developmental stage.

**Results:** We observed relationships between the expression of genes involved in *voltage-gated cation channel activity* during Early Midfetal, Adolescence and Early Adulthood timepoints and association with schizophrenia and bipolar disorder, such that genes more strongly associated with these disorders had relatively low expression during Early Midfetal development and higher expression during Adolescence and Early Adulthood. The relationship with schizophrenia was strongest for the subset of genes related to calcium channel activity, whilst for bipolar disorder the relationship was distributed between calcium and potassium channel activity genes.

**Conclusions:** Our results indicate periods during development when biological pathways related to the activity of calcium and potassium channels may be most vulnerable to the effects of genetic variants conferring risk to psychiatric disorders. Furthermore, they indicate key time points and potential targets for disorder-specific therapeutic interventions.

## Introduction

Biological insight to psychiatric disorders has come from the identification of molecular pathways enriched for genetic association, determined by large cohort genome-wide studies. However, the expression of many genes varies as a function of age, and therefore the relevance of genetically associated pathways likely varies across development.

Schizophrenia, bipolar disorder (BD) and major depressive disorder (MDD) show considerable heritability and share a substantial component of genetic risk (1–6). Variance between the disorders may in part be attributed to differences in the degree of neurodevelopmental impairment (3, 7, 8). Many psychiatric symptoms first present during adolescence or in early adulthood (9), implying that pathophysiology emerges as the brain matures (10). However, there is substantial evidence that altered neurodevelopment during earlier prenatal or postnatal periods may contribute to some, if not all, psychiatric disorders (3, 11–14). Identifying the developmental stage at which particular biological pathways are most likely to contribute to risk for psychiatric disorders is therefore an important step towards understanding disease etiology and targeting new treatments.

Genetic association studies of schizophrenia have consistently demonstrated a convergence of genetic risk upon sets of genes with synaptic functions (15–19), including discrete signal transduction pathways such as voltage-gated calcium channel complexes and glutamate receptor complexes. The most recent genome-wide association studies (GWAS) of BD and MDD also highlight genes and pathways related to synaptic activity (20–23). Hence, signalling complexes in the synaptic membrane represent strong candidates for psychiatric drug targeting.

Neurodevelopment is regulated by a program of tightly controlled gene expression (24, 25). The majority of GWAS loci linked to psychiatric disorders are non-coding and most likely mediate risk by affecting gene expression (26–29). Previous evidence highlights that genes harbouring risk for schizophrenia and BD are also among those most dynamically expressed across development, particularly in prefrontal cortical regions (30–33). However, the question of the developmental stage at which genes and gene pathways associated with major psychiatric diseases are primarily expressed remains largely unresolved, limiting our ability to reduce risk and target treatments.

By integrating developmental transcriptomic data of the human brain with GWAS data of genetic risk in schizophrenia, BD and MDD, we aimed to identify time points at which gene sets implicated in risk for psychiatric disorder are most strongly expressed, in view of highlighting periods when key biological pathways may preferentially confer risk to these disorders.

## Methods and Materials

### Genotype data

We obtained common variant summary statistics for schizophrenia, BD and MDD from published meta-analyses of genome-wide association study (GWAS) data. The schizophrenia sample (40,675 cases, 64,643 controls) (17) is a meta-analysis of a GWAS derived from UK cohorts of patients taking clozapine (CLOZUK; 11,260 cases, 24,542 controls) and an international Psychiatric Genomics Consortium (PGC) study (29,415 cases, 40,101 controls) (34). Case-control samples for BD were compiled by the PGC from 32 cohorts of European descent (20,352 cases, 31,358 controls) (21). Genotype data for MDD (135,458 cases and 344,901 controls) were derived from a PGC meta-analysis of seven cohorts of European descent (22).

### Transcriptomic data

Human brain transcriptomic data from across the lifespan were obtained from two sources. A primary dataset derived from post-mortem dorsolateral prefrontal cortex (DLPFC) of 336 individuals with no history of psychiatric condition, referred to as BrainSeq (31). Samples ranged in age from second trimester to 85 years (Supplementary Table 1). Tissue acquisition and processing are described previously (35). Total RNA was extracted from DLPFC grey matter and poly(A)-captured libraries were sequenced using Illumina HiSeq 2000 (31). Samples were also genotyped, as described (31), to allow for control of genetic ancestry. To check for replication across datasets, we obtained a second, smaller human developmental transcriptomic dataset from the Allen Institute BrainSpan Atlas (36, 37). We retained sequencing data from DLPFC samples (N = 40) ranging from the first trimester to 40 years (Supplementary Table 1). 16 samples from BrainSeq and BrainSpan originated from the same individuals; these were removed from BrainSeq to ensure independence of the datasets.

Raw sequencing reads from Brainseq and BrainSpan were processed using the same software pipeline. We applied FastQC (38) and Trimmomatic (39) to quality check and trim poor quality or technical sequences as appropriate. Reads were mapped to hg38 reference genome using HISAT2 (40) and genomic features were annotated using featureCounts (41). Gene counts were converted to RPKM values (calculated in relation to the number of gene-assigned reads).

### Gene sets

Gene sets defined by biological pathways were curated from the Gene Ontology (GO) database (42) (January 13^th^ 2020), excluding gene annotations with evidence codes NAS (Non-traceable Author Statement), IEA (Inferred from Electronic Annotation) or RCA (inferred from Reviewed Computational Analysis). For primary analyses, gene sets containing fewer than 100 genes were excluded to minimize the effect of outliers. After filtering, 1766 gene sets remained (Supplementary Table 2).

### Gene set association analysis

Gene set association analysis was performed in MAGMA v1.07 (43). Using GWAS summary statistics, SNPs with a minor allele frequency greater than 1% were annotated to genes. A window of 35 kb upstream and 10 kb downstream of each gene was allowed to include proximal regulatory regions (44). Gene-wide association statistics were calculated using the SNP-wise mean model with the 1000 Genomes European reference panel (45) to control for linkage disequilibrium. Gene set association was calculated in a one-tailed competitive test, using a background of all genes and conditional on the brain expression of each gene. Brain expression was defined as log_2_(RPKM + 0.5) where RPKM is the mean average across BrainSeq samples (excluding BrainSpan samples).

Primary analyses aimed to identify gene sets significantly enriched for GWAS association signal. Following correction for multiple testing using the false discovery rate (FDR) (46), stepwise conditional analyses were applied to significantly (FDR < 0.05) associated gene sets in order to identify a reduced number of independent sets that efficiently summarise the biological themes underlying the association enrichment. During this process, gene sets were repeatedly re-tested for genetic association, each time selecting the gene set with the highest effect size (Beta) and adding that set to the conditional variables. On each iteration, gene sets that were no longer nominally significant (unadjusted *P* ≥ 0.05) were excluded.

In secondary analyses, we refined the associated genes sets by partitioning them into subsets based on function or expression and testing the relative association of these to the larger set. To do this, we added the larger set to the model as a conditional variable. Comparisons of genetic association between non-overlapping gene sets were performed using a z-test of beta values.

### Developmental gene expression scores

Transcriptomic data were divided into developmental stages (Supplementary Table 1). If fewer than two samples were available for a developmental stage in BrainSpan or BrainSeq, that stage was not analysed and the samples excluded from that dataset. Accordingly, two samples from BrainSeq were excluded (one Early Fetal, one Late Infancy). Late Adulthood was not represented in BrainSpan. To permit comparison between genes and control for covariates, for each gene we constructed a score measuring the expression of that gene at each developmental stage relative to all other stages (30). Expression scores were calculated by fitting a linear regression model to each gene using limma (v.3.40.6) (47) and voom (48) and extracting the t-statistic for each developmental stage term:

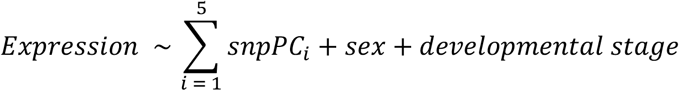

Developmental stage is a binary variable indicating whether the sample is from that stage or not. We controlled for genetic ancestry by covarying for the first five principal components defining sample genotypes (snpPC).

### Expression-association relationships

Within each independent gene set showing significant association enrichment to a disorder, we assessed the relationship between gene expression during a particular developmental stage and association with that disorder using MAGMA (v1.07) interaction analysis (49):

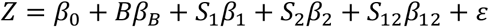

In this model, the interaction term *S*_12_ is defined as the product of gene set membership *S*_1_ and gene property *S*_2_ terms. This method determines whether enrichment for genetic association in a gene set is stronger for genes that have a higher gene property (in this case gene expression score). Due to the increased sensitivity of interaction analyses to outlier effects in smaller gene sets, primary analyses were restricted to sets containing at least 100 genes (49). Consistent with the gene set association analyses described previously, we used a background of all genes and conditioned on the mean brain expression of each gene. Interaction analyses were two-tailed. We corrected for the total number of tests performed on each gene set at each developmental stage using the Bonferroni method to give adjusted *P*-values (*P*.adj). To enable comparison across disorders, analyses yielding significant interactions in one disorder were repeated as secondary analyses in the other two disorders.

Since gene expression scores are calculated relative to the expression of the same gene at other developmental stages, this method enabled us to identify timepoints when genetically associated genes are more preferentially expressed.

### Clustering of developmental expression trajectories

To generalise expression analyses specific to each developmental stage to the complete expression trajectory across all stages, we partitioned gene sets of interest into subsets of genes with similar trajectories. Unexpressed genes (0 RPKM in all samples) were excluded from analysis. For each gene in a given gene set, expression at each developmental stage (mean RPKM) was scaled by subtracting the mean and dividing by the standard deviation. Clustering was performed using K-means. A suitable number of clusters was identified using a scree plot and clustering was run using 20 sets of random partitions and allowing for a maximum of 20 iterations. Gene set association analyses were used to test for enrichment of GWAS signal in the expression trajectory clusters. Conditional analyses were used to test for additional enrichment for association compared to the full set.

## Results

### Independently associated gene ontologies

To select gene sets for downstream analyses of developmental expression patterns, we performed gene set association analysis on a filtered set of 1766 GO-derived gene sets, as described above. Following correction for FDR (alpha = 0.05), we observed 62 gene sets significantly enriched for association with schizophrenia (Supplementary Table 2) and 17 associated with BD (Supplementary Table 3). No gene sets remained significantly associated with MDD after multiple testing correction (Supplementary Table 4). Due to the redundancy between GO terms, we used conditional analyses to select gene sets with independent associations. This procedure yielded 8 gene sets independently associated with schizophrenia (Supplementary Table 2) and 4 independently associated with BD (Supplementary Table 3). These analyses highlighted genes related to *voltage-gated cation channel activity* (*VG-cation*) as enriched for association with schizophrenia (β = 0.46, FDR = 5.5×10^−04^) and BD (β = 0.34, FDR = 0.011), but not MDD (β = 0.14, FDR = 1.0).

### Developmental stage-specific relationships between gene expression and genetic association within biological pathways

We used BrainSeq transcriptomic data to determine whether the enrichment for common variant association in gene sets with significant independent main effects is stronger for genes preferentially expressed during particular developmental stages, compared to a background of all genes. In analyses of genes annotated by *VG-cation*, we observed a significant positive interaction term for the relationship between expression during Early Adulthood and genetic association with both schizophrenia (β = 0.17, *P*.adj = 0.0030; Supplementary Table 5) and BD (β = 0.17, *P*.adj = 5.7×10^−04^; Supplementary Table 6) (Figure 1A,B). This indicates that, during Early Adulthood, *VG-cation* genes more strongly associated with schizophrenia or BD have relatively high DLPFC expression compared to other developmental stages, whilst those lacking association have relatively low expression. Conversely, during Early Midfetal development (10-15 post-conceptual weeks), there was a significant negative relationship (indicating that genes more associated with the disorder have relatively low DLPFC expression and less associated genes have higher expression) between *VG-cation* expression and association with BD (β = −0.052, *P*.adj = 0.0064) and evidence for the same relationship in schizophrenia, albeit non-significant (β = −0.055, *P*.adj = 0.053).

**Figure 1.**
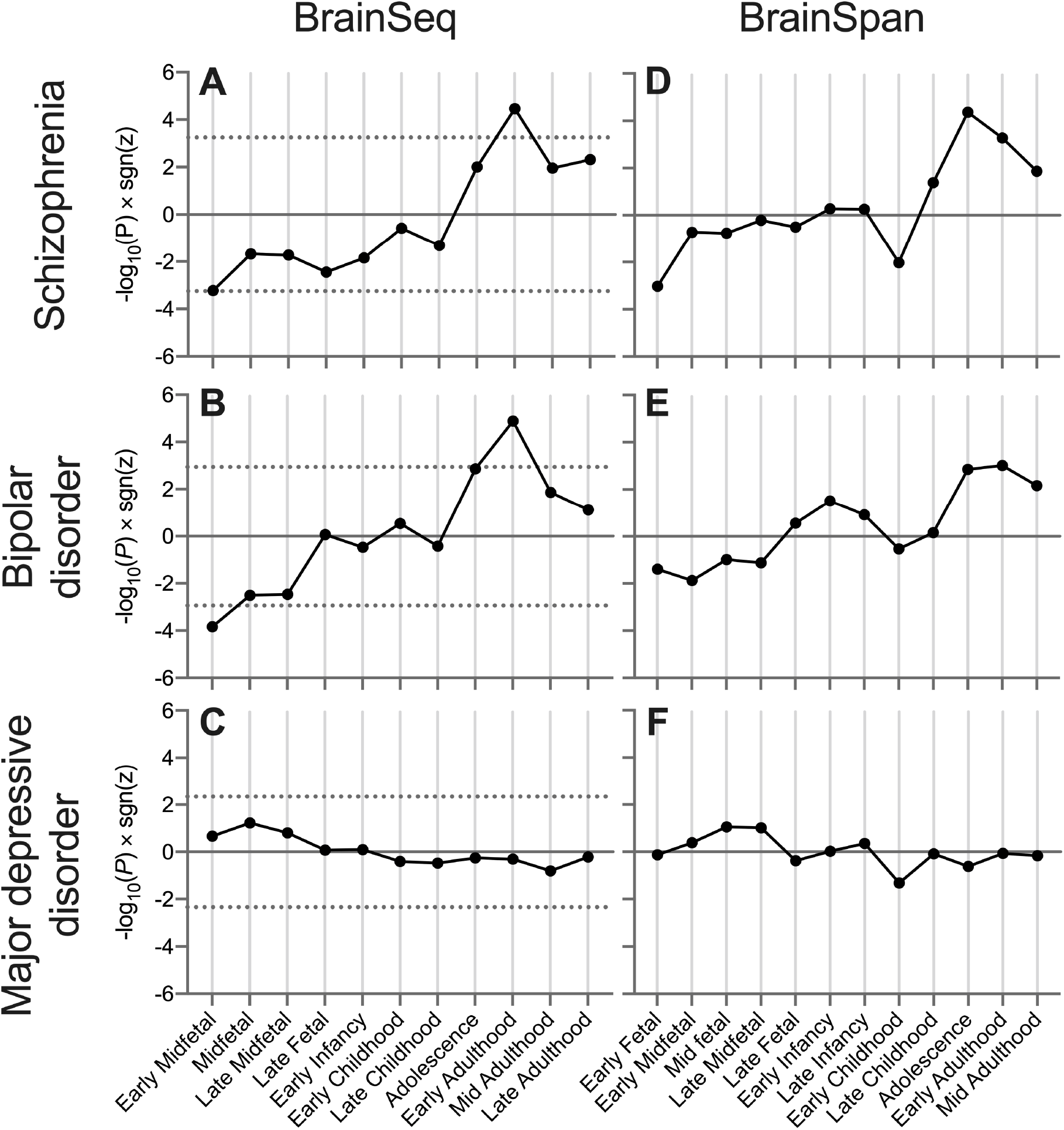
Developmental stage-specific relationships between the expression of *voltage gated cation channel activity* genes and genetic association with psychiatric disorders. Shown is −log_10_(*P*)×sgn(*z*) from independent MAGMA interaction analyses of common variant association and gene expression scores derived from BrainSeq or BrainSpan at each developmental stage, compared to a background of all genes. Dotted lines indicate thresholds for statistical significance after correcting for analyses of all gene sets with independent association with the disorder, and all stages (Supplementary Figures 5,6). MDD analyses were corrected for the number of stages only. Developmental stages containing fewer than 2 samples in the dataset are not represented. Early Fetal, 8-9 postconceptual weeks (pcw); Early Midfetal, 10-15 pcw; Midfetal, 16 pcw; Late Midfetal, 17-23 pcw; Late Fetal, 24-37 pcw; Early Infancy, 0-5 months; Late Infancy, 6-11 months; Early Childhood, 1-5 years; Late Childhood, 6-12 years; Adolescence, 13-19 years; Early Adulthood, 20-29 years; Mid Adulthood, 30-59 years; Late Adulthood, 60-100 years.

For comparison, we tested these temporal relationships with *VG-cation* expression in MDD genetic risk. *VG-cation* expression was not related to association with MDD at any developmental stage (Figure 1C).

We assessed the reliability of our findings by repeating the analyses of developmental *VG-cation* expression using an independent RNA sequencing-derived transcriptomic dataset, BrainSpan. Overall, the pattern of preferential *VG-cation* expression across development was consistent between BrainSeq and BrainSpan (Figure 1D-F), such that early embryonic expression exhibited the strongest negative relationship with association with schizophrenia (Early Fetal: β = −0.19, *P* = 9.4×10^−04^; Early Midfetal: β = −0.071, *P* = 0.19) and BD (Early Fetal: β = −0.10, *P* = 0.040; Early Midfetal: β = −0.11, *P* = 0.013), whilst strong positive relationships were observed during Early Adulthood (schizophrenia: β = 0.37, *P* = 5.3×10^−04^; BD: β = 0.32, *P* = 9.6×10^−04^). BrainSpan *VG-cation* gene expression during Adolescence was also strongly positively related to association with schizophrenia (β = 0.35, *P* = 4.3×10^−05^) and BD (β = 0.23, *P* = 0.0014). Again, no relationships were observed between *VG-cation* temporal expression and MDD association.

### Partitioning association between child terms of voltage-gated cation channel activity

The *VG-cation* term (N = 125 genes) is made up of four nested “child” GO terms: *voltage-gated potassium channel activity* (*VG-potassium*, N = 80), *voltage-gated calcium channel activity* (*VG-calcium;* N = 38), *NMDA glutamate receptor activity* (N = 7) and *voltage-gated proton channel activity* (N = 1). To test if genetic association of *VG-cation* with schizophrenia and BD is further enriched in the smaller and more biologically specific child terms, we performed a gene set analysis of the child terms (excluding *voltage-gated proton channel activity* due to small gene set size) conditional on *VG-cation* (Table 1). We observed that *VG-calcium* showed greater enrichment for association with schizophrenia (conditional *P* = 2.4×10^−05^) than the full *VG-cation* set. Conversely, association with BD was not further enriched within the child terms *VG-calcium* (conditional *P* = 0.20), *VG-potassium* (conditional *P* = 0.91) or *NMDA glutamate receptor activity* (conditional *P* = 0.15), indicating even distribution across subtypes of *VG-cation*.

**Table 1.** The relative association of *voltage-gated cation channel activity* child terms compared to the complete gene set, across two psychiatric disorders. Shown is the output from gene set association analysis in MAGMA before (*P*) and after (Cond *P*) conditioning on the full *voltage-gated cation channel activity* gene set.

| Gene set | N | Schizophrenia |  |  | Bipolar disorder |  |  |
| --- | --- | --- | --- | --- | --- | --- | --- |
| | | $\beta$ | $P$ | Cond $P$ | $\beta$ | $P$ | Cond $P$ |
| Voltage-gated cation channel activity | 125 | 0.46 | $9.3 \times 10^{-07}$ | NA | 0.34 | $4.1 \times 10^{-05}$ | NA |
| Voltage-gated calcium channel activity | 38 | 1.07 | $9.8 \times 10^{-10}$ | $2.4 \times 10^{-5}$ | 0.46 | 0.0029 | 0.20 |
| Voltage-gated potassium channel activity | 80 | 0.12 | 0.17 | 1.0 | 0.26 | 0.0070 | 0.91 |
| NMDA glutamate receptor activity | 7 | 1.11 | 0.0050 | 0.062 | 0.71 | 0.026 | 0.15 |

We examined the contribution of child terms *VG-calcium* and *VG-potassium* to the developmental patterns of preferential risk gene expression in *VG-cation* by performing interaction analyses on genes from either of these terms. We observed that the developmental relationships during Early Midfetal and Early Adulthood stages between gene expression and association with schizophrenia were more prominent in *VG-calcium* genes than *VG-potassium* genes (Early Midfetal: z-test *P* = 0.027; Early Adulthood: z-test *P* = 0.013) (Figure 2A). Conversely, these same relationships in analyses of BD appeared more pronounced among *VG-potassium* genes (Figure 2B), although comparisons of effect sizes showed no significant differences at these stages (Early Midfetal: z-test *P* = 0.63; Early Adulthood: z-test *P* = 0.21). This is likely due to the larger gene set size of *VG-potassium* compared to *VG-calcium* giving a greater significance for a given effect size. These analyses also strengthened the evidence that these positive expression-association relationships begin in Adolescence and continue into Early Adulthood.

**Figure 2.**
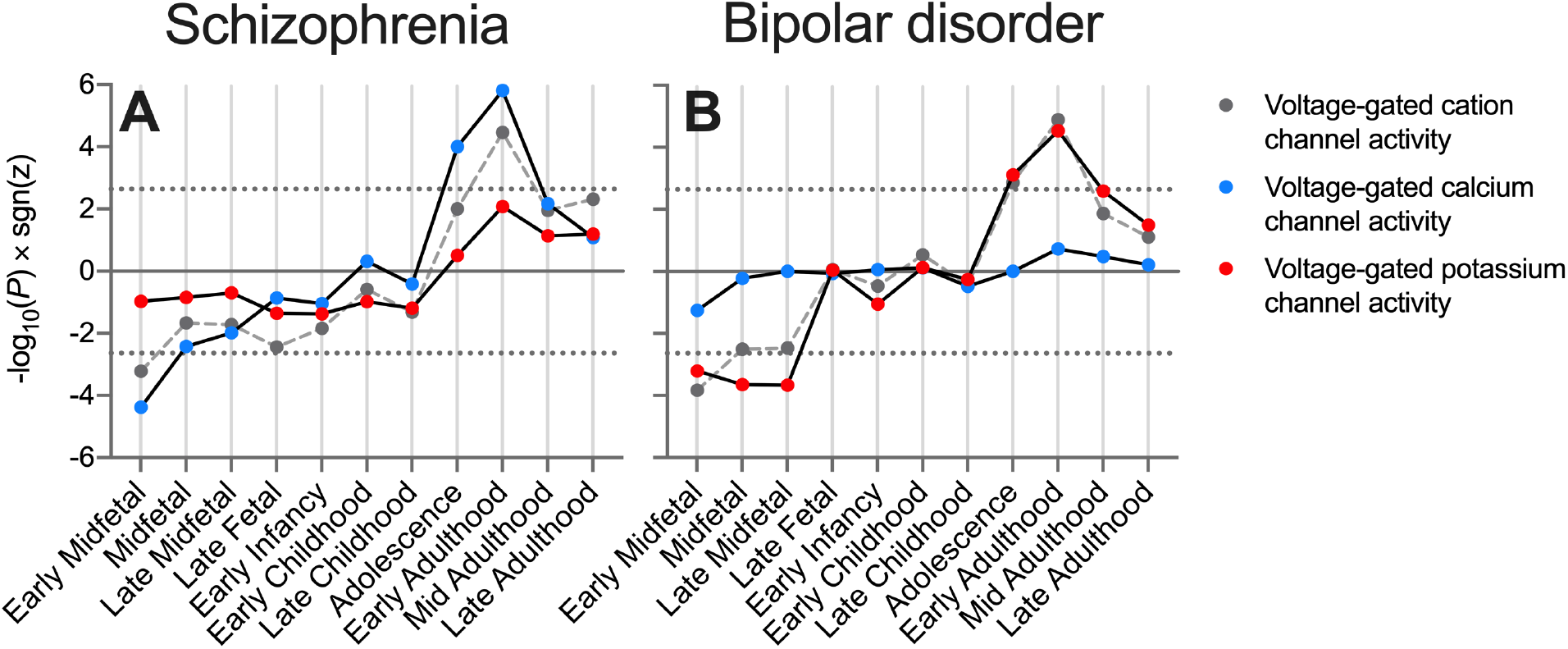
Comparison of expression-association relationships at 11 developmental stages within *voltage-gated cation channel activity* child terms. Shown is −log_10_(*P*)×sgn(*z*) from MAGMA interaction analyses of schizophrenia or BD common variant association and BrainSeq-derived gene expression scores at each stage, compared to a background of all genes. Dotted lines indicate thresholds for statistical significance after correcting for analyses of two gene sets across 11 stages. GO:0022843 *Voltage-gated cation channel activity*, N = 125; GO:0005245 *Voltage-gated calcium channel activity*, N = 38; GO:0005249 *Voltage-gated potassium channel activity*, N = 80.

Together, these results suggest that within the set of genes responsible for regulating the activity of voltage-gated cation channels, themselves strongly enriched for genetic association with schizophrenia and BD, the subset involved in calcium activity show further enrichment for association with schizophrenia, with those more strongly associated being preferentially expressed during Adolescence and Early Adulthood. On the other hand, the calcium and potassium subsets exhibit a more balanced association with BD, and the Early Adulthood preferential expression of BD associated genes involved in calcium activity is no greater than (or even less than) that of potassium activity genes.

### Partitioning association by developmental expression trajectory

Our analyses of developmental stage-specific expression suggest that *VG-cation* genes with low Early Midfetal and high Adolescence / Early Adulthood expression are enriched for association with schizophrenia and BD. They also suggest that such an enrichment in schizophrenia is stronger for the *VG-calcium* subset. To generalise these hypotheses across all developmental stages we used clustering to identify subsets of genes showing similar expression trajectories and performed gene set association analyses to test the distribution of genetic association. We subsequently repeated these analyses conditioning on the full *VG-cation* set to compare the relative enrichment of association. K-means clustering identified four broad expression trajectories in *VG-cation* genes (Supplementary Figure 2; Supplementary Table 7; Figure 3). Enrichment for genetic association with schizophrenia and BD was restricted to Clusters 1 and 2, both of which contain genes with lower embryonic expression and higher expression in later life (Figure 3). However, of the two subsets, only Cluster 2 (containing genes exhibiting later peak expression than Cluster 1) harbored greater enrichment for association with schizophrenia and BD than *VG-cation* genes as a whole (schizophrenia: conditional *P* = 1.9×10^−05^; BD: conditional *P* = 0.0037).

**Figure 3.**
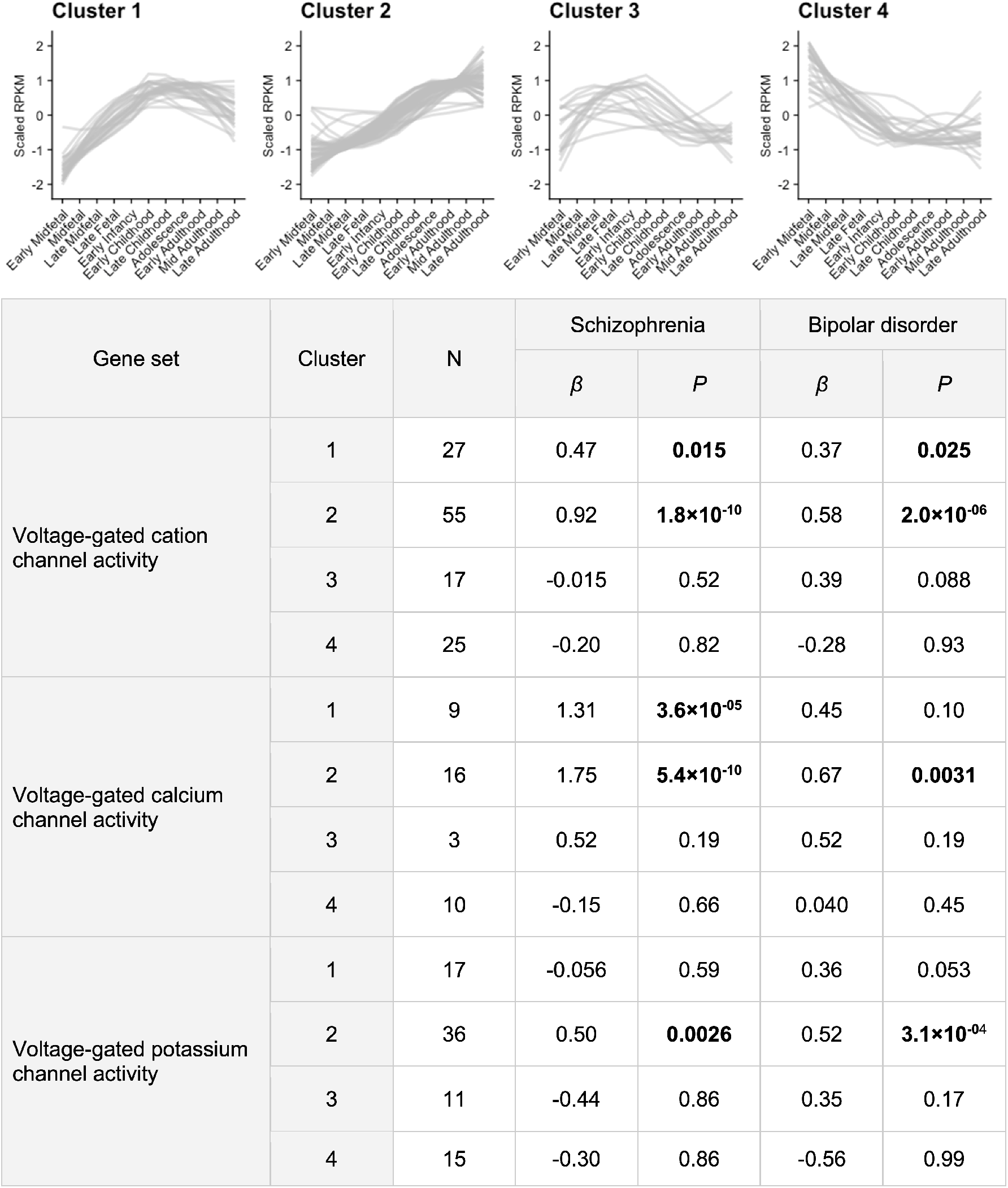
Clusters of developmental expression trajectories of *voltage-gated cation channel activity* genes and their enrichments for GWAS association in schizophrenia and BD. K-means clustering was used to divide genes based on their mean RPKM expression in BrainSeq across 11 developmental stages. Tabulated are results from schizophrenia and BD gene set association analyses of each expression trajectory cluster, with separation into *voltage-gated calcium channel activity* and *voltage-gated potassium channel activity* child terms.

On splitting these clusters into *VG-calcium* and *VG-potassium*, we observed that association of Clusters 1 and 2 with schizophrenia was stronger among *VG-calcium* genes, such that the effect size was greater than *VG-potassium* genes (Cluster 1: z-test *P* = 9.5×10^−04^; Cluster 2: z-test *P* = 2.2×10^−04^) and only the *VG-calcium* subsets showed greater enrichment for association than *VG-cation* as a whole (*VG-calcium* Cluster 1: conditional *P* = 0.0038; *VG-calcium* Cluster 2: conditional *P* = 8.1×10^−07^; *VG-potassium* Cluster 1: conditional *P* = 0.99; *VG-potassium* Cluster 2: conditional *P* = 0.44). Conversely, association with BD of *VG-cation* genes with the same expression trajectories was distributed between *VG-calcium* and *VG-potassium* subsets (Cluster 1: z-test *P* = 0.82; Cluster 2: z-test *P* = 0.60). Neither subset of Cluster 1 or Cluster 2 exhibited significantly greater enrichment for association with BD than the full *VG-cation* gene set (*VG-calcium* Cluster 1: conditional *P* = 0.36; *VG-calcium* Cluster 2: conditional *P* = 0.068; *VG-potassium* Cluster 1: conditional *P* = 0.47; *VG-potassium* Cluster 2: conditional *P* = 0.071).

Hence, the schizophrenia common variant association of genes responsible for regulating the activity of voltage-gated cation channels is most enriched in the subset involved in calcium activity characterised by an upward developmental expression trajectory in the DLPFC, peaking in adolescence or later. This is in contrast with BD association, which is enriched in genes with the same expression trajectory, but with no skew towards either cation type.

## Discussion

Our study highlights the biological convergence among common variants conferring risk to schizophrenia and BD in genes annotated by the ontological term *voltage-gated cation channel activity* (*VG-cation*) and identifies adolescence and early adulthood as periods of preferential expression for the most strongly associated members in both disorders. We demonstrated that in schizophrenia these temporal relationships derived predominantly from the calcium-related component of *VG-cation*, *voltage-gated calcium channel activity* (*VG-calcium*), whilst in BD they were driven by both *VG-cation* and *voltage-gated potassium channel activity* (*VG-potassium*) components. Genetic association with MDD was enriched in *VG-calcium* genes, but we observed no relationship between this association and preferential gene expression at any developmental stage.

An enrichment for genetic association in genes encoding voltage-gated calcium channel subunits has been reported in primary GWAS of schizophrenia, BD and MDD, on which this study was based (17, 21, 22). There has been consistent evidence supporting their involvement in these psychiatric disorders from common variant studies (34) and in schizophrenia from rare variants (19, 50). The voltage-gated calcium channel family (consisting of L, T, P/Q, N and R subtypes) are signal transducers of electrical excitation in neurons. Through the induction of intracellular calcium transients, they couple membrane depolarization to signalling cascades which include activity-dependent regulation of gene expression (51). In this way, voltage-gated calcium channels influence synaptic plasticity and are thought to be important drivers of learning and memory (52).

We observed that genetic association with BD in *VG-cation* was distributed between calcium and potassium components. Voltage-gated potassium channels have been linked to BD through genetic studies previously (21, 53–57), and this represents a divergence from schizophrenia and MDD which have not shown evidence of conferring risk through potassium channels. The broad function of voltage-gated potassium channels is to control the threshold for action potentials and repolarise the membrane after firing (58). Like calcium channels, they have been shown to contribute to synaptic plasticity (59, 60).

It is notable that the *VG-cation* GO term examined here does not include subsets related to the activity of sodium channels. Voltage-gated sodium channels have been linked to schizophrenia and autism in sequencing studies (50, 61, 62) and are involved in the pharmacodynamics of several mood stabilizers used in the treatment of bipolar disorder, including valproic acid (63). Whilst the current study found no significant enrichment for association with schizophrenia, BD or MDD among GO terms *sodium ion transport* or *sodium ion transmembrane transporter activity* (Supplementary Tables 2-4), the developmental profile of sodium channels associated through other types of genetic variant would benefit from further investigation.

Our results suggest that Adolescence and Early Adulthood are periods of peak preferential DLPFC expression of *VG-cation* genes associated with schizophrenia and BD. This lends to the idea that these developmental stages, which correspond to peak periods of symptom onset, may be more vulnerable to risk conferred through cation channels. Conversely, Early Midfetal development (corresponding to 10-15 post-conceptual weeks) was characterised by comparably low expression of risk *VG-cation* genes, perhaps indicating a period of low vulnerability in this pathway. We cannot rule out, however, that variation of function of a gene expressed in relatively low abundance could mediate substantial risk.

More specifically, these periods of preferential gene expression in schizophrenia-associated *VG-cation* genes were characterised by more prominent relationships within the *VG-calcium* subset. Conversely, preferential expression of BD-associated *VG-cation* genes was agnostic to the cation subset. This discrepancy raises the possibility that only *VG-calcium* genes contribute to an Adolescence / Early Adulthood vulnerability to schizophrenia, whilst both *VG-calcium* and *VG-potassium* genes contribute to BD vulnerability during the same time period.

Whilst our results reflect trends within gene sets, the expression patterns of individual genes encoding cation channel subunits enriched for association with schizophrenia or BD may vary. Furthermore, each gene may encode multiple transcripts with differing time periods of peak expression. For example, *CACNA1C* (encoding calcium channel CaV1.2) contains a locus with genome-wide evidence for association with schizophrenia, BD and MDD (17, 21, 64–66) and is the single-gene cause of Timothy syndrome, a disorder of multiorgan maldevelopment (67). *CACNA1C* is reported to reach peak brain expression in late fetal / early childhood development (68, 69) and its forebrain deletion in embryonic, but not adult, mice models endophenotypes of psychiatric disorders (70). However, around 250 splice variants of *CACNA1C* have been reported (71). These transcripts were shown to exhibit varying regional expression profiles and are predicted to vary in their developmental regulation as well. Further, it is still unclear which of the *CACNA1C* transcripts is associated with risk variants identified from GWAS. To better understand developmental vulnerability to genetic risk conferred through cation channels, the expression patterns of transcripts relevant to psychiatric disorders should be characterised independently of the gene-wide expression.

Our study employed transcriptomic data derived from the DLPFC, a region long thought to be impacted in schizophrenia and mood disorders (72–74). Compared to other brain regions, the DLPFC is considered to be late-maturing (75, 76) and exhibits some distinct developmental gene expression trajectories (77, 78). Hence the relationships between genetic risk and developmental expression in other brain regions might be different or create age-specific vulnerabilities in other biological pathways, perhaps accounting for the emergence of different psychiatric symptoms over time.

Beyond levels of expression, there are undoubtedly other factors influencing vulnerability to genetic risk across the lifetime, untested in this study. These may include activity of compensatory mechanisms, development-specific functions or susceptibility to environmental insult, among others. Moreover, there is strong evidence that early developmental stages contribute to psychiatric pathophysiology, yet manifest as a disorder later in the lifetime as the brain matures (11, 73). Using similar methodology, our previous work (30) demonstrated that among all brain-expressed genes there is a bias for those with stronger genetic association to be preferentially expressed in the prefrontal cortex during early midfetal development (schizophrenia) or early infancy (schizophrenia / BD). Therefore, it is very likely that other biological pathways enriched for association with psychiatric disorders are most vulnerable to genetic risk during prenatal or early postnatal life, but were not identified by the current study.

Our findings have clear implications for the targeting of therapeutics. Psychiatric disorders are insufficiently treated, yet voltage-gated cation channels have been recognised as potential targets for new (or existing) compounds (60, 79) and their druggability is an area of active research (80, 81). Preferential expression of genetically associated genes during ages typical of diagnosis reinforces the suitability of voltage-gated cation channels as targets, and suggests that different targets may be particularly relevant in schizophrenia and BD. Our results also indicate periods when therapeutic agents acting on such pathways may be most effective.

## Acknowledgements

This study was supported by Medical Research Council (MRC) grants MR/L010305/1 and G0800509 and a Wellcome Trust Strategic Award (100202/Z/12/Z). We thank the Bipolar Disorder and Major Depressive Disorder workgroups of the Psychiatric Genomics Consortium for providing summary statistics used in this study. We would like to thank the research participants and employees of 23andMe for making this work possible.

We thank R. Zielke, R. D. Vigorito and R. M. Johnson of the National Institute of Child Health and Human Development Brain and Tissue Bank for Developmental Disorders at the University of Maryland for providing fetal, child and adolescent brain specimens. We also acknowledge the following brain bank collections: the Mount Sinai NIH Brain and Tissue Repository, the University of Pennsylvania Alzheimer’s Disease Core Center, the University of Pittsburgh NeuroBioBank and Brain and Tissue Repositories and the NIMH Human Brain Collection Core. CMC leadership: Pamela Sklar and Joseph Buxbaum (Icahn School of Medicine at Mount Sinai), Bernie Devlin and David Lewis (University of Pittsburgh), Raquel Gur and Chang-Gyu Hahn (University of Pennsylvania), Keisuke Hirai and Hiroyoshi Toyoshiba (Takeda Pharmaceuticals Company Limited), Enrico Domenici and Laurent Essioux (F. Hoffman-La Roche Ltd), Lara Mangravite and Mette Peters (Sage Bionetworks), and Thomas Lehner and Barbara Lipska (NIMH).

