## Supplementary Information for "Developmental profile of psychiatric risk associated with voltage-gated cation channel activity"

**Supplementary Figure 1** Comparisons of the relationship between GWAS association with schizophrenia or bipolar disorder and the relative expression of *voltage-gated cation channel activity* genes in BrainSeq during Early Midfetal and Early Adulthood developmental stages. Gene expression scores were scaled by subtracting the mean and dividing the standard deviation. Genetic association is represented by log converted gene-wide *P*-values from MAGMA analysis using the SNP-wise mean model, controlling for linkage disequilibrium.
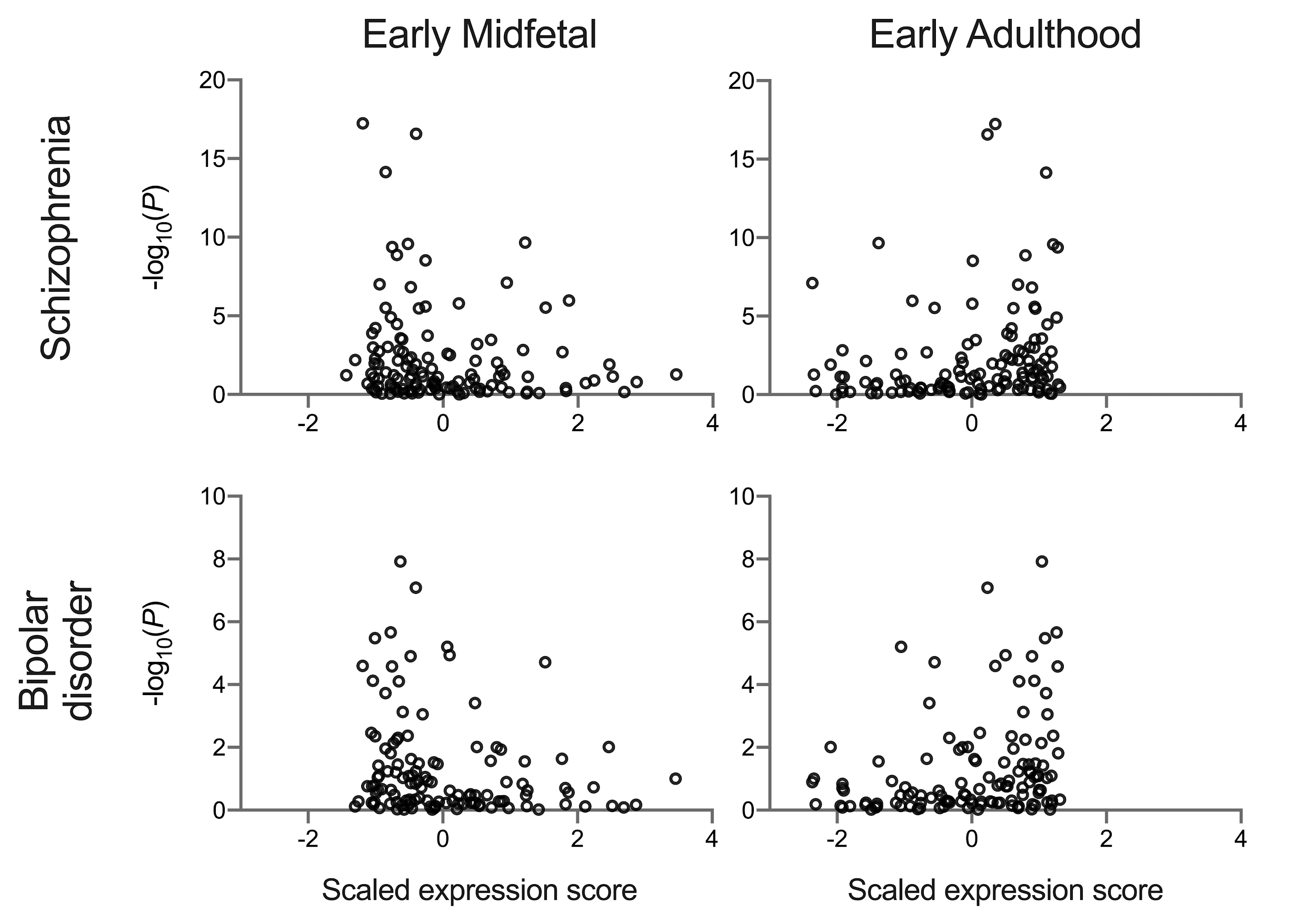

**Supplementary Figure 2** Scree plot from K-means clustering of *voltage-gated cation channel genes*. Clustering was performed for 1:15 centroids. Each run used 20 sets of random partitions, allowing for a maximum of 20 iterations per run.
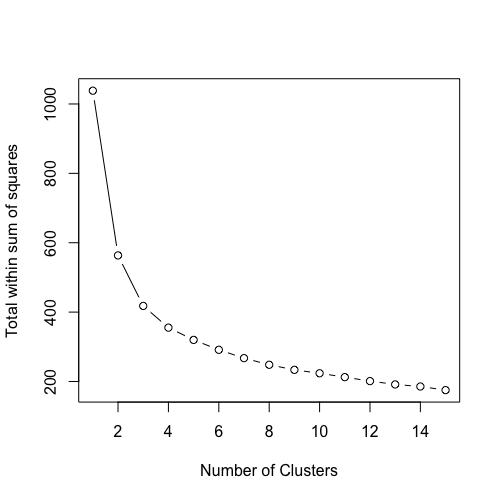
